# 3D mapping of disease in ant societies reveals a strategy of a specialized parasite

**DOI:** 10.1101/003574

**Authors:** Raquel G. Loreto, Simon L. Elliot, Mayara L. R. Freitas, Thairine M. Pereira, David P. Hughes

**Author notes:** Corresponding authors: Raquel G Loreto. W241A Millennium Science Complex, Pennsylvania State University, Pennsylvania State University, University Park, PA 16802, USA. David Hughes. W129 Millennium Science Complex, Penn State University, Pennsylvania State University, University Park, PA 16802, USA.

## Abstract

Despite the widely held position that the social insects have evolved effective ways to limit infectious disease spread, many pathogens and parasites do attack insect societies. Maintaining a disease-free nest environment is an important evolutionary feature, but since workers have to leave the nest to forage they are routinely exposed to disease. Here we show that despite effective social immunity, in which workers act collectively to reduce disease inside the nest, 100% of studied ant colonies of *Camponotus rufipes* in a Brazilian Rainforest were infected by the specialized fungal parasite *Ophiocordyceps unilateralis s.l*. Not only is disease present for all colonies but long-term dynamics over 20 months revealed disease is a permanent feature. Using 3D maps, we showed the parasite optimizes its transmission by controlling workers’ behavior to die on the doorstep of the colony, where susceptible foragers are predictable in time and space. Therefore, despite social immunity, specialized diseases of ants have evolved effective strategies to exploit insect societies.

## Introduction

High density living in human settlements, or among the animals and plants we raise for food can result in both major epidemics and the emergence of novel pathogens [1]. Dense societies also occur in natural systems and the prime example are the social insects. Their colonies can contain thousands and sometimes millions of highly related individuals [2] which might suggest constant epidemics. However, such societies have in fact achieved both ecological dominance and long evolutionary stability. For instance, ants have become dominant in terrestrial biomes accounting for over 50% of the biomass despite making up less than 2% of all insect species [3]. This success implies that their societies have evolved ways to reduce disease pressure, a phenomenon known as social immunity [4]. Therefore, important lessons for limiting disease spread might be gained from examining societies that have evolved by the process of natural selection over long periods of time.

Many studies have demonstrated that ants, termites, bees and wasps successfully defend their colonies from a range of parasites, through an integration of different levels of immunity, from cellular to behavioral [4–7]. However, this raises a paradox since we know that many different groups of parasites do infect social insects and, based upon their life history, the majority appear to be specialized parasites requiring infection of colony members for lifecycle completion [6–8]. If colonies are so adept at defending the nest, an important question is: How do specialized parasites transmit despite such effective group defense?

The majority of studies on disease dynamics in ants have focused their attention on colony response to generalist parasites introduced inside the nest, under laboratory conditions [5,9–11]. These studies can control numerous factors and capture quantitative details of collective defensive strategies. However, such studies minimize environmental complexity, missing important components of the host parasite interaction. For example, they do not focus on the ants that are more exposed to infection (foragers) and do not consider specialized parasites, which have adaptive traits that drive transmission. Without incorporation these factors, they do not explain if and how co-evolved parasites infect their host, somehow circumventing the social immunity. Thus, we set out to study a disease of ants in a rainforest ecosystem, incorporating environmental complexity.

Further we addressed specialization by focusing on a parasite that has evolved the ability to control host behavior to affect transmission [12–16] and is specific to its host species [14].

## Material and Methods

### Study area and host and parasite species

Fieldwork was carried out at the Research Station of Mata do Paraíso, Universidade Federal de Viçosa, Minas Gerais, Southeast Brazil (20°48′08 S 42°51′31 W). The carpenter ant *Camponotus rufipes* is very abundant in this habitat. The ants forage on trails, being active at night with activity peaking in the early evening [17]. The trails are built mainly on twigs and branches lying on the forest floor so the ants use the 3D space of the forest, not walking only on the floor. Ant trails are permanent with the same trail being used for weeks [17]. The entomopathogenic fungi *Ophiocordyceps unilateralis sensu lato* is a specialized parasite of ants that must kill the host to complete it life cycle. Before killing the infected ant, the fungal parasite manipulates the behavior of host, leading the hosts to climb the vegetation, bite the veins and margins of leaves in rainforests [12–16], that then serve as a platform for fungal growth and spore release from the dead ant [15]. *Ophiocordyceps unilateralis s.l*. transmission requires the growth of a long stalk from the head of a dead ant from which spores are released onto the forest floor to infect other workers. The newly described parasitic fungus *Ophiocordyceps camponoti-rufipedis*, previously known as *O. unilateralis s.l*. [14], is a parasite that has *C. rufipes* as its host, and it is also abundant in the study area [17].

### Disease within the nest

Experimentally it has been shown that the fungus cannot grow either on the forest floor or in the dry upper canopy [12], thus the manipulation is adaptive for the parasite. While informative, these earlier experiments left open the question of if the parasite could develop inside the nest. To determine whether the fungus is able to grow normally inside an ant mound, we collected a whole nest of *C. rufipes* (including nest material, ants and brood) and a recently abandoned nest (only the nest material, no ants or brood). Both were directly placed in buckets (volume = 8L), maintaining the original characteristics of the nest, and kept under natural day/night light and temperature regime. We collected 28 ants freshly killed by *Ophiocordyceps camponoti-rufipedis*, took pictures of their initial conditions and attached them to flags (so as not to lose them inside the nest). Each cadaver was at the fungal pre-emergence stage, meaning the ant’s body had been colonized by fungal blastospores and hyphae but the stalk required for transmission had not yet grown. Since the stalk is crucial for transmission we identified this stage as being crucial for fungal lifecycle completion. The 28 ants were placed in two different treatments: (1) nest with ants, n=14; (2) nest without ants, n=14. In the treatment with ants, sugar/water 50% and canned tuna were used to feed the ants. In both treatments the fungal killed ants were placed on the top of 10 cm of nest material and covered with 20 cm of the same nest material. The ant cadavers were removed 10 days later, and pictures were taken to evaluate the development of the fungus.

### Disease surrounding the nest

Because social immunity is well known from experimental studies in the laboratory to be effective and rapidly deployed [5,9–11] we might expect colonies in nature to be disease-free. We therefore set out to ask how common infection by *O. camponoti-rufipedis* was at the population level. In order to identify nests of *C. rufipes*, we made 22 transects of 2,000m^2^ each (100x20m), covering 44,000m^2^. The first 15 transects were initiated on the main path of the research station, and were taken 100m into the forest, using string as a guide reference. From the string, 10m for both sides were covered. In order to obtain more complete coverage of the site, we delineated a new path from which we traced the other six transects, covering the 2,000m^2^ area for each one. The distance between the start points of each transect was 100m and the transect direction alternated between the left and right sides of the path. Using this methodology we found 9 nests. Another 8 previously identified nests were used in this study. We examined the vegetation within a 1m radius around each nest looking for ants killed by the O. *camponoti-rufipedis*, attached to the underside of leaves. The nests that had dead ants on the adjacent vicinity were recorded.

To investigate the disease surrounding the nest with more details and over longer periods of time, we mapped the dead ants surrounding 4 nests over 20 months (Dec 2010-Jul 2012). Because the ants are known to travel long distances from their nests [18], we limited the mapping to the area surrounding the nest. This area is critical because the ants must walk throughout there when leaving or returning to the nest. We also refer to this area as “doorstep” of the colony. For 4 of the 17 previously discovered nests, we demarcated a study area of 200m^3^ (10m x 10m x 2m) (from now called plot) that were centered on the nest. Thus, we observed the long-term dynamic of the fungal infection in four distinct colonies.

In order to determine the 3D position of ants killed inside the four studied plots, we used the coordinate system relative to the nest, determining the x, y and z position of each dead ant, having the left bottom corner as beginning. For example, all the four nests had the coordinates (500, 500, 0) because they were in the center of the plots (x=500cm, y=500cm) and on the forest floor (z=0cm). We measured the disease in 3D (x,y,z coordinates) because the ants are manipulated to die attached to the underside of leaves on plants in the understory vegetation of tropical forests [12–16].

Before beginning the 3D measures, in November 2010 we tagged all dead ants in those plots checking every single leaf inside the plots, up to 2m from the forest floor. Across the first six consecutive months (December 2010-May 2011) we identified, tagged and mapped (x,y,z coordinates) every single newly killed ant attached to leaves within each of the plots. None of the dead ants that we counted were removed from the plots so we did not reduce the naturally occurring disease pressure. To capture long-term dynamics, we left the area for seven months following the May 2011 census and mapped the new cadavers in January 2012. Finally, we returned in July 2012 to check if each of the four nests had new dead ants on the immediate vicinity of the colony.

Since the potential hosts are encountered on the foraging trails, we also measured and mapped in 3D the trails formed by the ant. The foraging trails were marked with small flags placed every 30cm, starting at the nest and continuing until they left the plot. The coordinates of each flag inside the area were determined the same way as we did for the dead ants. The z positions, measured from the forest floor, were included because the trails pass along on branches, lianas and roots above the forest floor [17]. Combining those coordinates we were able to access the exact location of each trail in space, related to the nest. We also did not disturb the ant trails. The trails and dead ants were mapped once a month, being necessary a day for each plot (trails and dead ants for each plot were mapped at the same day). The 3D data were plotted using the Grasshopper® plugin for the 3D modeling platform Rhino®. Statistical analyses were conducted using R (version 2.15.2). We used generalized mixed models to avoid temporal pseudo-replication, using the variable “Month” as repeated factor.

## Results

### Disease within the nest

Of the total of 28 samples placed inside the nest, none developed normally (Fig. S1). Eight (53%) of the cadavers placed within the nest material without ants did not grow at all, and the remaining six (47%) grew abnormally in a way that ensured spore transmission would not occur, since it occurs from a specialized structure (ascoma) that grows near the top of that stalk that grows from the ant’s head) (Fig. S1 B, D). Of the cadavers placed into nests containing live ants, nine were removed from the leaf they were attached to (64%) and it was not possible to find them. This suggests that the healthy ants removed the cadavers, possibly destroying them since we could not recover them. The remaining five (36%) failed to grow normally meaning, as occurred in the nest without ants, that sporulation did not happen and hence no transmission occurred. In summary, the fungal parasite was incapable of reaching the infective stage inside ant nests, whether ants were present or not.

### Disease surrounding the nest

We discovered 17 nests that were patchily distributed in the study area. All 17 nests had ant cadavers attached to leaves beside the ant colony. Thus, disease prevalence at the population level is 100%.

For the long-term study, during these first six consecutive months (Dec 2010-May 2011), we identified 347 newly dead ants, killed by *O. camponoti-rufipedis* surrounding the four colonies (Fig. 1, Movie S1). The number of dead ants is month-dependent (Mixed-model: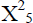: 60.877; P<0.0001). December 2010 had the highest density of parasitized ants: 146 dead ants attached to leaves were found in the census for that month (Mixed-model: 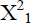: 18.052; P<0.0001) (Fig. 2). The lowest occurrence of dead ants was in March 2011, when we recorded a total of 12 dead ants; but it did not differ statistically from February (24 dead ants) (Mixed-model: 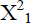: 2.0164; P=0.1556) (Fig. 2). November, December and January receive 75% of the yearly precipitation [19], which is likely an important determinant of abundance for fungal parasites. In January 2012, after we left the plots for seven months, we found a total of 39 new dead ants within the 4 plots (that is, after seven months, each nest had freshly killed cadavers attached to leaves). Finally, when we returned in July 2012 and established that, even after 20 months, each of those four nests had ants manipulated to die in the immediate vicinity of both the nest and trails, demonstrating the long-term persistence of disease surrounding these colonies.

**Figure 1.**
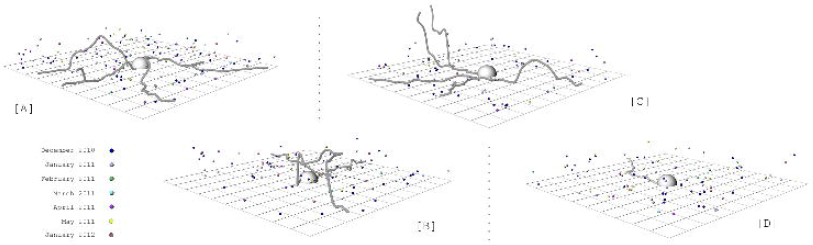
3D maps of foraging trail and monthly-infected ants surrounding ant colonies in Atlantic rainforest, Brazil. The infected ants represent accumulated dead ants in 7 months (Dec 2010-May2011 and Jan 2012). Distinct colors represent different months. The lines show trails were recorded in December 2010. (A) Nest A. (B) Nest B. (C) Nest C. (D) Nest D.

Over the 20-month period we measured disease in eight months (i.e. months 1-6, 7, and 20) surrounding four colonies. Only once and for one colony we did not find new records of *O. camponoti-rufipedis* surrounding host colonies (Nest C) (Fig. 2). This was the month that the density of new cadavers was lowest for all colonies (March 2011) (Fig. 2). However, in the following month (April 2011), we did find newly manipulated killed ants outside Nest C, demonstrating that the colony was not disease free.

**Figure 2.**
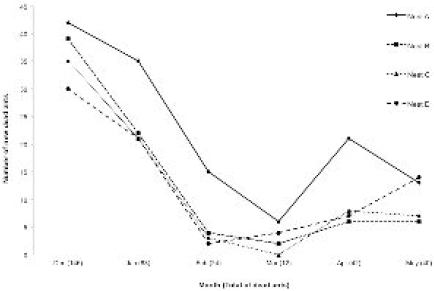
Disease dynamics surrounding four nests of *Camponotus rufipes* in an Atlantic rainforest fragment, Brazil. Different lines represent each analyzed nest (A, B, C or D). The numbers show the total of dead ants recorded for each of the six months.

Because we measured and mapped the position and abundance of ant trails, we also investigated the role of host activity on the disease dynamics. We calculated the number of trails for each month, which represent the healthy ants susceptible to new infections, and related it with the number of dead ants of each month. We would expect to find more dead ants when the ants were more active (activity was measured by number of trails). Surprisingly, the number of infected ants attached to leaves surrounding the colony was not related with the number of susceptible hosts (Mixed-model: 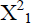: 2.1078; P=0.1466).

## Discussion

Our results support the previously well-established theory of social immunity operating inside the nest of social insects, as we have shown that the *Camponotus rufipes* ants removed most of the *Ophiocordyceps camponoti-rufipedis* parasitized cadavers placed within the nest. Additionally, however, we also found that this specialized fungal parasite, when placed inside a nest without ants, cannot grow to the stage suitable for transmission. Previous studies on disease in ant societies have shown that corpses are removed by nestmates [20–22] and sick ants experience social isolation [9,23]. Both of these behaviors are interpreted as a class of behavioral immunity that prevents diseases spreading among nestmates [9,20,23]. However, we showed that simply being within the nest reduces the fitness of the specialized parasitic fungus *O. camponoti-rufipedis* to zero, whether the nest is inhabited by ants or not. It may be that the removal of corpses and, more importantly, dying in social isolation (outside the nest) actually increases the opportunity for the parasite to complete its development and be transmitted to the next host. From the perspective of the colony, the ability of nestmates to destroy cadavers before the fungus can become infectious means that remaining inside the nest might better serve the colony compared to ants dying in social isolation outside the nest where fungal growth can occur. The same may apply for generalist pathogens, such as *Metarhizium* (used in the majority of studies on social immunity), that have a broad range of hosts [24]. These do not necessarily need to be transmitted from ant to ant and can be transmitted to others insect groups outside the nest.

Not all parasites of ants use the same strategy of manipulating the host as *Ophiocordyceps* that we studied. To place our results within the wider context of parasites evolved to infect ant societies we examined the mode of transmission for other specialized parasites of ants (Table S1). It was striking that, as with *Ophiocordcyeps*, the majority of parasites of ants require the infected ant to leave the nest to continue the life cycle. Social isolation mediated by parasites may be a widespread strategy in parasites that attack ant societies. These parasites only can only be within the nest when they are invisible to the nestmates, internal to the infected ant body. The life stage that requires them to either exit from or protrude from the host body occurs outside the nest, where the social immunity does not act.

Although social immunity is present in insect societies such as the ants studied here, and does function to prevent disease transmission within the nest, our full appreciation of it may not be wholly realized because to date we have been biased by studies that have focused solely on ant behavior towards diseases inside the nest. But as we showed and as is reflected in the literature (Table S1) most parasites of ants transmit outside the nest. To date, studies of social immunity have not considered the importance of the environment outside the nest for transmission and diseases dynamics. By mapping the disease *in natura*, we were able to graphically simplify the environment without reducing or eliminating any component of the system, making it possible to study disease dynamics for an insect society. In what is the first survey of a specialized disease of an ant colony in a rainforest we established that disease is a permanent feature (across 20 months) and it is present in 100% of the examined colonies (17 total).

### The terminal host model of transmission

Social immunity is effective and prevents disease transmission within the nest. From a host-centric view this would appear to provide an advantage to the host within the assumed arms race between the two parties. However, we offer an additional viewpoint. The fungus *O. camponotini-rufipedis*, infects susceptible hosts (foraging ant workers) by means of large curved spores that fall directly down from the cadavers attached to leaves that will be picked up by a new host [14].

Foraging for food is an indispensable task for the colony and workers must perform this task. Typically, it is a very risky job carried out by older workers, which are going to die sooner [25, 26]. In this scenario, where the older ants collect the food supplies required by the colony, there is a constant turnover of new susceptible ants on trails. We expect that the proportion of infected ants within a colony is low, since not all workers forage. As we have shown, the foragers are constantly being killed by the parasite, and new workers will take over the risky tasks that are done outside the nest, providing a continual stream of new hosts for the parasite that sits right on the doorstep of the colony. Probably over the long term such a strategy has impact on the host demography and social interactions, although evidence remains lacking on this important question.

Targeting a specific group within a population or group of cells within a body is a widespread strategy in antagonistic interactions. For instance, many predators attack weak prey, which include old, sick and young individuals. These are easier to capture as they occupy peripheral positions on outside of the herd or simply lag behind in chases and because of their weak status are undefended. Considering within-body host-parasite interactions the papillomavirus uses the strategy of a high reproductive rate in terminal cells, which is considered advantageous because there is no immune surveillance in such cells [27]. This virus, which transmitted by contact, forms warts on the most external surface of the host body - skin, enabling the transmission to a new potential host [27]. The trematoda *Euhaplorchis californienis*, a killifish parasite, has also evolved the strategy of making its host occupy external positions within a group: by changing host swimming behavior, the parasites increases the probability of predation which is advantageous as the parasite reaches it final host – the avian predator [28]. We suggest that the specialized parasite *O. camponotini-rufipedis* specifically targets older individuals from ant societies and causes them to die on the doorstep of the colony. The advantage is that the parasite does not need to evolve mechanism to overcome the effective social immunity that occurs inside the nest, and at the same time, it ensures a constant supply of susceptible hosts.

An option for the host would be extending the social immunity to the outside nest environment. There are anecdotal observations of ants removing fungal manipulated and killed cadavers from the environment. In the Amazon rainforest, the turtle ant, *Cephalotes atratus*, which is arboreal, removes the cadavers from the bark of trees [29].

The wood ant, *Formica rufa*, which inhabits grasslands, remove the cadavers on the doorstep of the colony, surrounding their nest [30]. It would be of great interest to test how far out from the colony social immunity can extend. In the 20 months of fieldwork we did not see any ants destroying or removing the cadavers attached to leaves surrounding the nest, leading us to suspect that they do not display the same defensive behavior around the nest as they do inside. It is likely difficult or costly for ants to control the outside environment, where *O. camponoti-rufipedis* is strategically positioned. Although social immunity does not occur outside the nest in this case, it might be possible for adaptive changes in ant behavior to reduce the disease burden. The species of ant we studied builds their foraging trails using bridges and this might function to reduce contact with the soil and establish the permanent use of the same pathway, both of which might decrease the risk of infection [17,31]. We do know examples of how the foraging trail network of ants adaptively shift in response to changing food abundance [32,33] or to reduce the incidence of attack by predators [34,35,36] or competing colonies [37,38]. There are also examples of trails shifting in presence of parasitoid females that lay eggs in workers [39–41]. Also, the presence of *O. unilateralis* in Thailand was suspected of causing the target ant, *Camponotus leonardi* to reduce the time spent near the forest floor [13]. Generally, ant trail behavior and its response to parasites are neglected but through our focus on within forest parasite-host dynamics we hope to encourage such work. However, because foraging ants tend to be older there may simply be little selection on the host to evolve strategies against the parasite. If this is the case then host and parasite may not be involved in a coevolutionary arms race (as is commonly assumed) at all. They may both be following quite stable evolutionary strategies.

### Conclusion

The concept of the social insect colony as a “factory constructed inside a fortress” [42] does present a challenge for parasites “breaking into the fortress” [6]. The highly evolved social immunity system is the first line of defense that until now appeared highly effective. Such was the supposed efficacy that early genome studies were quick to point out honeybees had 1/3 the immune genes of other insect because behavior was considered so important, obviating the need for humoral immunity [43]. But the existence of many specialized parasites of ant societies (Table S1) demonstrates parasites can and do transmit despite this collective defense. Our focus on a co-evolved specialized parasite in a complex tropical rainforest environment has highlighted a weakness in what is an otherwise effective barrier: workers need to leave the confines of the nest to collect food. This means susceptible hosts are predictable in both time and space and parasites have evolved to exploit this. As we emphasized throughout, the view that insect societies, such as ants, are paragons of effective collective defense against disease transmission [44] has largely been developed from studies conducted under artificial conditions. We have shown that within the complex arena of a rainforest, specialized diseases of ants have evolved effective methods to constantly transmit to new hosts by controlling worker ants to die on the doorstep of the colony. Since the majority of specialized parasites do require a transmission step outside the colony (Table S1), we would expect they also exploit the vulnerability of colonies where workers must leave to forage. Taken all the results together this implies that while social immunity is effective within the nest, it does not function against specialized parasites because they have evolved strategies to transmit outside the nest, consequently not encountering the social immunity.

## Acknowledgements

RGL was initially funded by CNPq and is currently supported by CAPES (#6203/10-8). SLE is funded by CNPq. DPH is funded by Penn State University. We are grateful to Camila Moreira, Diana Goodman and Lauren Quevillon for assistance in fieldwork and Andrew Read, Christina Grozinger and Jim Marden for the discussion.

## Supplementary Figures Legend

**Figure S1: *Camponotus rufipes* ants infected infected by *Ophiocordyceps camponoti-rufipedis*.** (A) Ant recently killed by the specialized parasite *Ophiocordyceps camponoti-rufipedis*. (B) Mature *O. camponoti-rufipedis* stage, suitable to transmission. The arrow points to the frutification body from where the spores are shot. (C) Collected ant recently killed by the fungus parasite before on the experiment. Fungal presents initial development (arrow). (D) Same sample after 10 days inside the host nest. The fungal did not developed as it normally does outside the next (arrow).

**Movie 1: Spatiotemporal dynamics of the specialized fungal parasite attacking an ant colony across six consecutive months.** The data was collected in Atlantic rainforest, southeastern Brazil. The funfal parasite species is *Ophiocordyceps camponoti-rufipedis* that attacks the ant host *Camponotus rufipes*. The red dots represent the new ants killed by the parasite in the respective month. The grey dots represent the sum of dead ants from previous months. The red lines represent the forage trail on the ant host for each studied month.

**Table S1: Overview of co-evolved parasites in ant societies.** Transmission can be between ants (direct) or also include another host (indirect). The final environment, where the sexual reproduction of the parasite occurs, can be in the environment surrounding the nest (Outside the nest), within the colony (Inside the nest) or final host 500 (Vertebrate host). The effect of parasitism is often death of the infected, either directly attributable to the parasite (Direct death), or indirectly via a behavioral change that leads to the host being eaten by the final host (Predation) or jumping in water, to allow the parasite to enter water for mating (Drowning). Additional details of each group in Schmid-Hempel (1998) [6].

